# Deficiency in ribosome biogenesis causes streptomycin resistance and impairs motility in *Salmonella*

**DOI:** 10.1101/2024.01.08.574728

**Authors:** Zhihui Lyu, Yunyi Ling, Ambro van Hoof, Jiqiang Ling

**Affiliations:** Department of Cell Biology and Molecular Genetics, The University of Maryland, College Park, MD 20742, USA; Department of Microbiology and Molecular Genetics, The University of Texas Health Science Center at Houston, Houston, TX 77030, USA

**Keywords:** Pathogen, antibiotic failure, ribosomal fidelity

## Abstract

The ribosome is the central hub for protein synthesis and the target of many antibiotics. Whereas the majority of ribosome-targeting antibiotics inhibit protein synthesis and are bacteriostatic, aminoglycosides promote protein mistranslation and are bactericidal. Understanding the resistance mechanisms of bacteria against aminoglycosides is not only vital for improving the efficacy of this critically important group of antibiotics but also crucial for studying the molecular basis of translational fidelity. In this work, we analyzed *Salmonella* mutants evolved in the presence of the aminoglycoside streptomycin (Str) and identified a novel gene *rimP* to be involved in Str resistance. RimP is a ribosome assembly factor critical for the maturation of the 30S small subunit that binds Str. Deficiency in RimP increases resistance against Str and facilitates the development of even higher resistance. Deleting *rimP* decreases mistranslation and cellular uptake of Str, and further impairs flagellar motility. Our work thus highlights a previously unknown mechanism of aminoglycoside resistance via defective ribosome assembly.

## Introduction

Antibiotic resistance has become an urgent threat to the global healthcare system. A recent study shows that bacterial antimicrobial resistance is associated with 4.95 million human deaths in 2019 (1). Aminoglycoside antibiotics have been historically used to treat infections caused by many pathogens, such as *Mycobacterium tuberculosis, Salmonella enterica*, and *Yersinia pestis*, and are currently listed as critically important antimicrobials for human therapy by the World Health Organization (2, 3). Aminoglycosides target the bacterial ribosomes and cause protein mistranslation (4, 5). It is proposed that mistranslated proteins primes aminoglycoside uptake in an energy-dependent process, leading to irreversible inhibition of the ribosome and cell death (6, 7).

Streptomycin (Str) is one of the first discovered and most extensively studied aminoglycosides (8-13). Structural analyses reveal that Str binds to the decoding center of the ribosomal small subunit comprising the 16S rRNA and ribosomal protein uS12 (RpsL) and stabilizes a closed conformation to promote codon-anticodon mismatch (13, 14). Mutations leading to Str resistance have been mapped to *rpsL, rrs*, and *rsmG* (*gidB*) genes (15-20). The *rrs* operon encodes rRNAs, whereas RsmG methylates G527 (*E. coli* numbering) of 16S rRNA near the decoding center (21, 22). These known mutations thus appear to directly affect the binding site of Str on the ribosome. In this study, we have identified *rimP* as a novel gene involved in Str resistance. RimP is a conserved bacterial protein contributing to the assembly of the 30S small ribosomal subunit (23, 24). We show that deleting *rimP* increases Str resistance 8-fold in both *Salmonella enterica* and *Escherichia coli*, and promotes evolution of higher resistance. The Δ*rimP* mutant also exhibits a lower rate of mistranslation in the presence of Str and a decreased level of Str uptake. Our work reveals that ribosome assembly impacts aminoglycoside action.

## Results

### Spontaneous Str-resistant mutants evolve rapidly in *Salmonella* culture

Bacteria acquire antibiotic resistance through horizontal gene transfer or spontaneous mutations (25). In animal hosts, Str-resistant *Salmonella* strains often arise through the acquisition of *strA* and *strB* genes (encoding Str-inactivating enzymes) from other bacterial species (26). In our study, we aim to explore the intrinsic mechanisms leading to Str resistance using *Salmonella enterica* Serovar Typhimurium as a model pathogen. We first performed evolution experiments with an increasing concentration of Str in Luria-Bertani (LB) broth. Wild-type (WT) *Salmonella* (ATCC 14028s) cells were inoculated in parallel experiments in LB with 0-128 μg/ml Str and grown to saturation. The cultures grown in the highest Str concentration were further inoculated in fresh media with Str. The WT *Salmonella* strain has a minimal inhibitory concentration of 16 μg/ml for Str (Fig. 1 and Table 1). After three rounds of evolution, all nine parallel lineages were able to grow in the presence of 128 μg/ml Str, indicating that Str-resistant strains evolve rapidly without horizontal transfer of foreign genes.

**TABLE 1.** Streptomycin, motility, and growth rates of *Salmonella* variants.

| Strain | Genotype | Str MIC (µg/ml) | Motility (mm) | Growth rate (h<sup>-1</sup>) |
| --- | --- | --- | --- | --- |
| WT | *S. Typhimurium* ATCC 14028s | 16 | 30.0 ± 2.4 | 2.56 ± 0.05 |
| M1 | *rsmG* (G159fs) | 2048 | 32.3 ± 2.7 | 2.46 ± 0.01 |
| M2 | *rsmG* (L168stop), *fre* (P55L) | 256 | 6.5 ± 2.6 | 2.22 ± 0.14 |
| M3 | *rsmG* (P118fs), *ΔispA* (partial) *ΔxseB*, *ΔthiI*, *ΔphnV*, *ΔphnU*, *ΔphnT*, *ΔphnS* | 4096 | 3.3 ± 0.5 | 2.56 ± 0.05 |
| M4 | *rsmG* (G156stop), *ubiD* (Q78stop) | 4096 | 3.2 ± 0.2 | 0.48 ± 0.05 |
| M5 | *ubiD* (Q78stop), *sbmA* (R165fs) | 512 | 3.5 ± 0.4 | 0.67 ± 0.07 |
| M6 | *ubiD* (Q78stop), *rimP* (G31fs) | 4096 | 3.3 ± 0.2 | 0.56 ± 0.007 |
| M7 | *rsmG* (R101W), *ubiD* (Q78stop) | 4096 | 3.2 ± 0.2 | 0.65 ± 0.07 |
| M8 | *rsmG* (R101L), *cyoB* (W203stop) | 512 | 12.8 ± 1.9 | 2.25 ± 0.09 |
| M9 | *rsmG* (T22fs), *aadA* promoter (G to A) | 512 | 30.8 ± 0.9 | 2.46 ± 0.14 |
| *ΔrimP* | *rimP*::Chl | 128 | 3.7 ± 0.4 | 1.78 ± 0.06 |
| *ΔrsmG* | *rsmG*::Chl | 128 | 26.5 ± 1.2 | 2.44 ± 0.03 |

**Figure 1.**
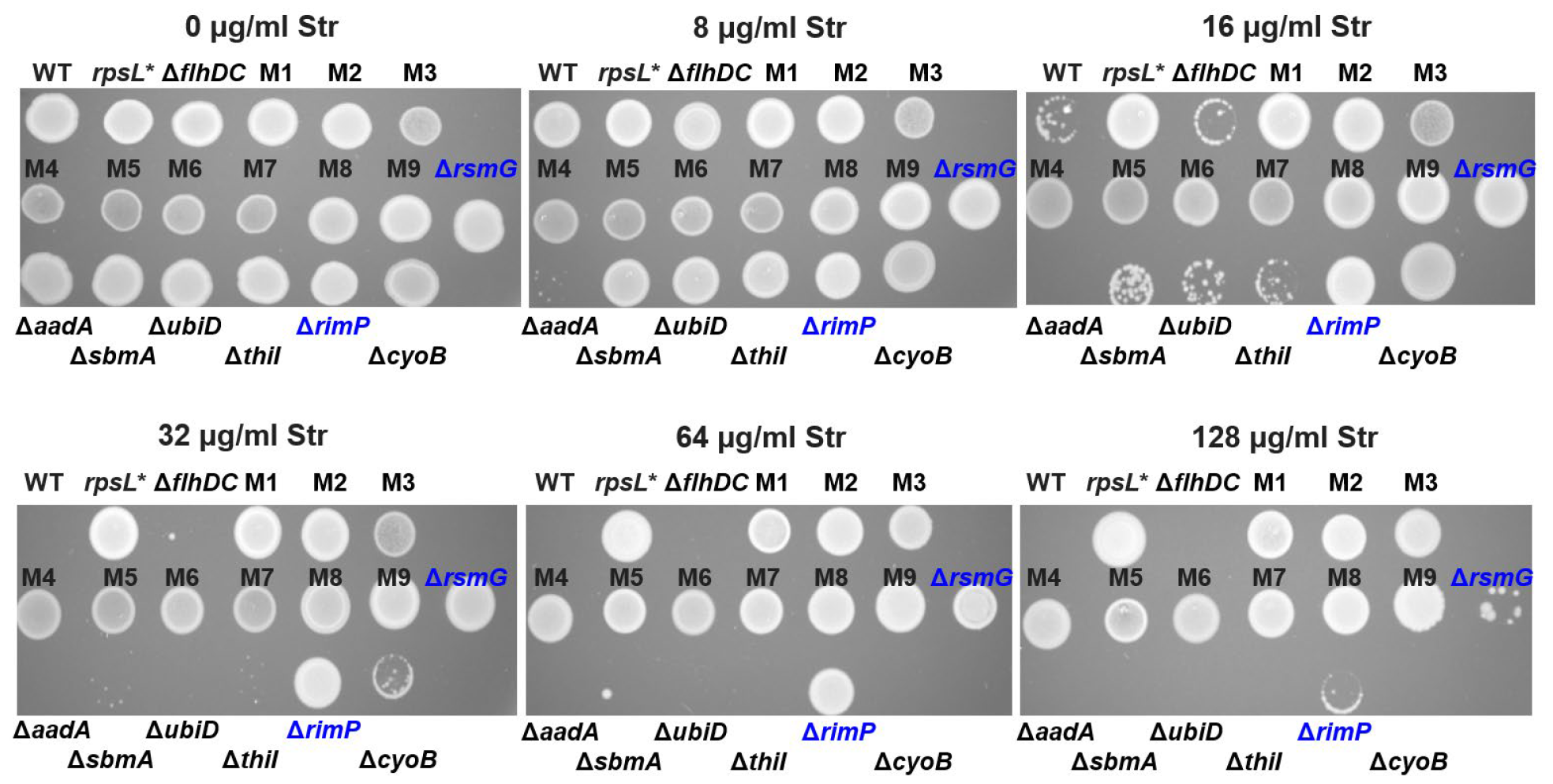
Streptomycin resistance in *Salmonella* Typhimurium variants. Overnight cultures of *Salmonella* variants grown in LB without Str were spotted on LB agar plates with various concentrations of Str. All the evolved strains (M1-M9) and *rpsL\** (carrying a K42N mutation in uS12) grew at 128 μg/ml Str. Deleting *flhDC*, the master regulator of flagellar genes, did not increase Str resistance. Deleting *rsmG* or *rimP* increased the MIC 8-fold. These images are representatives of at least three biological replicates.

### Mutations in *rimP* and *rsmG* promote Str resistance

We next performed whole-genome sequencing in an effort to identify the mutations responsible for Str-resistance in the evolved strains. Mutations in Str-resistant strains most frequently occur in *rsmG*, and are also found in *aadA, sbmA, ubiD*, the *phn* operon (including *thiI*), *rimP*, and *cyoB* (Table 1). To validate the mutations contributing to Str resistance, we constructed knock-out strains of the above genes. Deleting *rimP* or *rsmG* increased the minimal inhibitory concentration (MIC) for Str 8-fold from 16 to 128 μg/ml (Fig. 1 and Table 1). Single deletion mutants of *aadA, sbmA, ubiD*, and *thiI* did not affect the MIC of Str, whereas deleting *cyoB* increased the MIC 2-fold. The higher Str resistance in the evolved strains might have resulted from polar effects, a combination of mutations, or unidentified mutations. To test whether mutations in *rimP* and *rsmG* promote evolution of high Str resistance, we plated WT, Δ*rimP*, and Δ*rsmG* cells on LB agar with or without 512 μg/ml Str. Compared with the WT, both the Δ*rimP* and Δ*rsmG* strains exhibited significantly more spontaneous mutants that were resistant to 512 μg/ml Str (Fig. 2). Mutations in *rsmG* have been frequently identified in Str-resistant *Mycobacteria* and *Salmonella* strains (15, 18, 27), yet to our knowledge, *rimP* has not been previously shown to affect aminoglycoside resistance. This prompts us to further investigate the role of *rimP* and ribosome biogenesis in Str action in cells.

**Figure 2.**
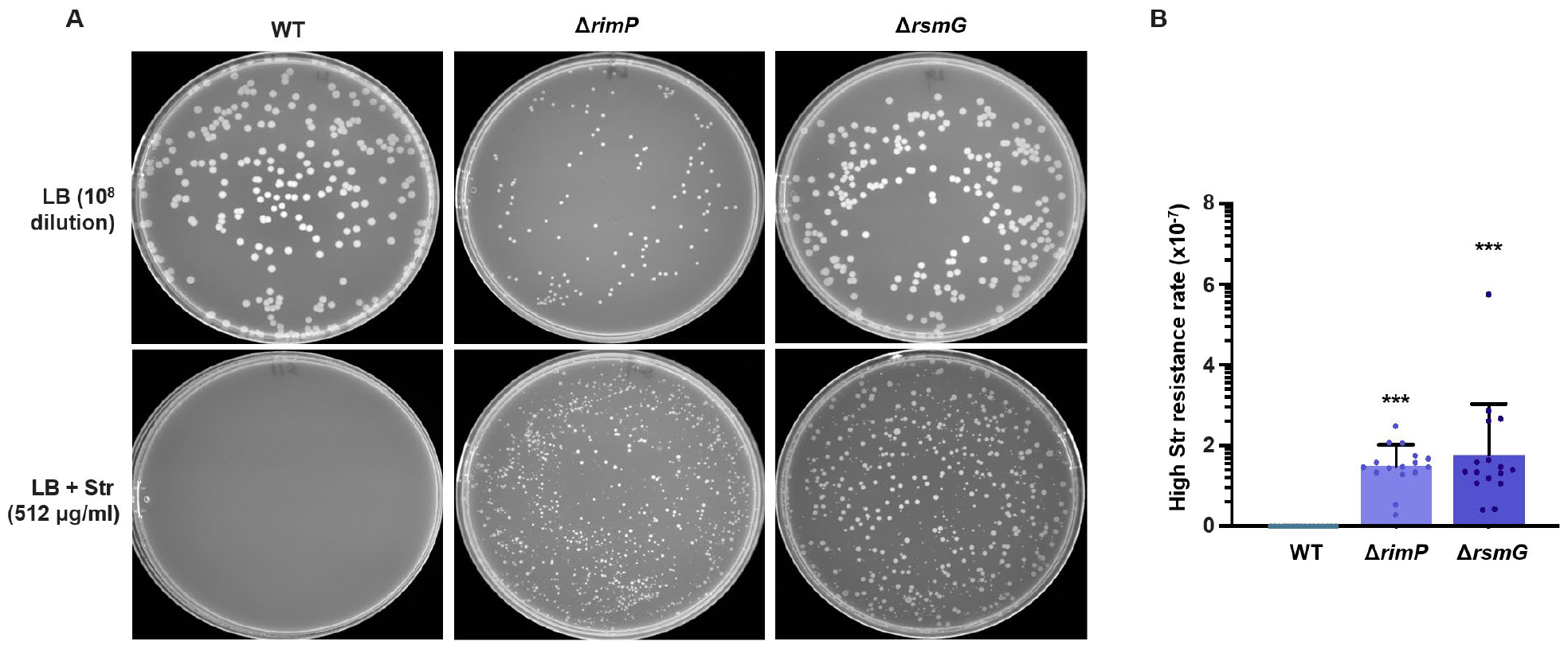
Fluctuation tests of Str resistance. (A) Representative images of colony formation on LB agar with or without Str. WT, Δ*rimP*, and Δ*rsmG Salmonella* were grown in LB at 37°C to the stationary phase and plated. (B) High Str resistance rates were calculated using the ratio of colony number on LB + Str over that on LB. Each dot represents one biological replicate. Error bars represent one SD from the mean. The statistical differences between the mutants and the WT were analyzed using One-way ANOVA with Dunnett correction. *** P < 0.001.

### RimP is critical for Str uptake

Previous studies suggest that mistranslated membrane proteins in the presence of aminoglycosides lead to membrane disruption and a faster phase of uptake (6, 7, 28). We used the fluorescent dye DiBAC_4_(3) (28, 29) to probe membrane potential and integrity of WT, Δ*rimP*, and Δ*rsmG* cells in the presence and absence of Str. As a control, we also included the *rpsL\** strain (carrying a K42N mutation in uS12) with high Str resistance (Fig. 1 and Table 1). The fluorescence signal of DiBAC_4_(3) is induced upon entry to the cytoplasm. Our flow cytometry analyses revealed that Str treatment substantially increased DiBAC_4_(3) fluorescence in WT cells, similar to treatment with the ionophore carbonyl cyanide m-chlorophenyl hydrazone (CCCP) (Fig. 3A). The fraction of DiBAC_4_(3)-positive cells in the Δ*rimP*, Δ*rsmG*, and *rpsL\** mutants upon Str treatment was much lower compared with the WT (Fig. 3A), suggesting that the membrane integrity was less disrupted by Str in these mutants. Further fluorescence microscopy and platereader analyses yielded results similar to flow cytometry (Figs. 3B and 3C). Next, we tested the relative concentrations of Str in WT and Δ*rimP* using a zone-inhibition assay as described in (30). The extracts of Str-treated *Salmonella* cells were spotted on LB agar plates with a lawn of Str-sensitive *E. coli*. We found that the Δ*rimP* extract displayed a smaller inhibition zone compared with the WT, indicating that deleting *rimP* lowered the intracellular concentration of Str.

**Figure 3.**
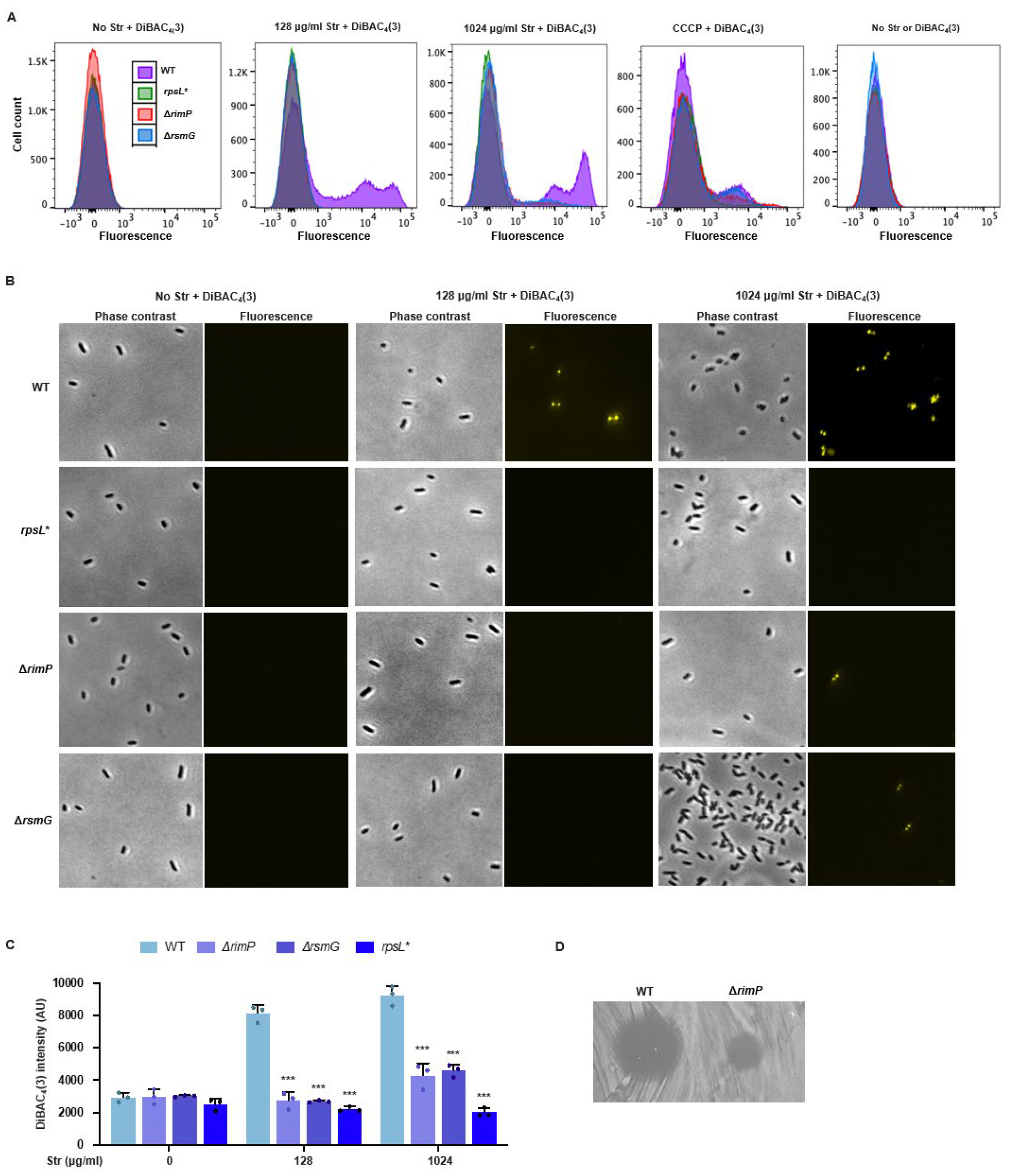
Membrane depolarization and Str uptake in *Salmonella* variants. Log-phase *Salmonella* cells were treated with or without Str at indicated concentrations and DiBAC_4_(3). The fluorescence was determined using (A) flow cytometry, (B) fluorescence microscopy, and (C) platereader. (D) Zone inhibition of the *E. coli* lawn by the lysates of WT and Δ*rimP Salmonella* treated with 1024 μg/ml Str. (A), (B), and (D) show representative images of at least three biological replicates. Error bars in (C) represent one SD from the mean. The statistical differences between the mutants and the WT at the same Str concentration were analyzed using One-way ANOVA with Dunnett correction. *** P < 0.001.

The intracellular concentrations of antibiotics are affected by uptake and efflux. A key component of efflux pumps in *Salmonella* and *E. coli* critical for multi-drug resistance is TolC (31). We found that deleting *tolC* in the WT *Salmonella* increased the sensitivity to Str (Fig. S1A), suggesting that TolC played a role in Str efflux. Compared with the Δ*tolC* mutant, the Δ*rimP*/Δ*tolC* double mutant was still more resistant to Str (Fig. S1A). We further used Nile Red to test the efflux activity. Whereas deleting *tolC* substantially increased intracellular accumulation of Nile Red due to an efflux defect, deleting *rimP* did not affect Nile Red efflux (Fig. S1B). In addition, deleting *rimP* did not increase resistance against tetracycline (Tet), ciprofloxacin (Cip), or ampicillin (Amp) (Fig. S2). Collectively, our data suggest that deleting *rimP* does not affect TolC-dependent efflux, but mainly decreases the uptake of Str.

### RimP enhances mistranslation in the presence of Str

Mutations in uS12 that lead to Str resistance often improve ribosomal fidelity, which counters the effect of Str to induce mistranslation (32, 33). Deleting *rsmG* also increases ribosomal fidelity in *E. coli* and *M. tuberculosis* (27, 34). Using a dual-fluorescence reporter that our lab previously developed (35, 36), we found that the *Salmonella* Δ*rsmG* strain exhibited decreased readthrough of UGA stop codons (Fig. 4), suggesting an improved ribosomal fidelity as in *E. coli* and *M. tuberculosis*. Deleting *rimP* slightly increased UGA readthrough in the absence of Str. However, in the presence of Str, the UGA readthrough level was significantly lower in the Δ*rimP* strain compared with the WT. Such lower mistranslation could explain the improved membrane integrity in the Δ*rimP* mutant upon exposure to Str (Fig. 3).

**Figure 4.**
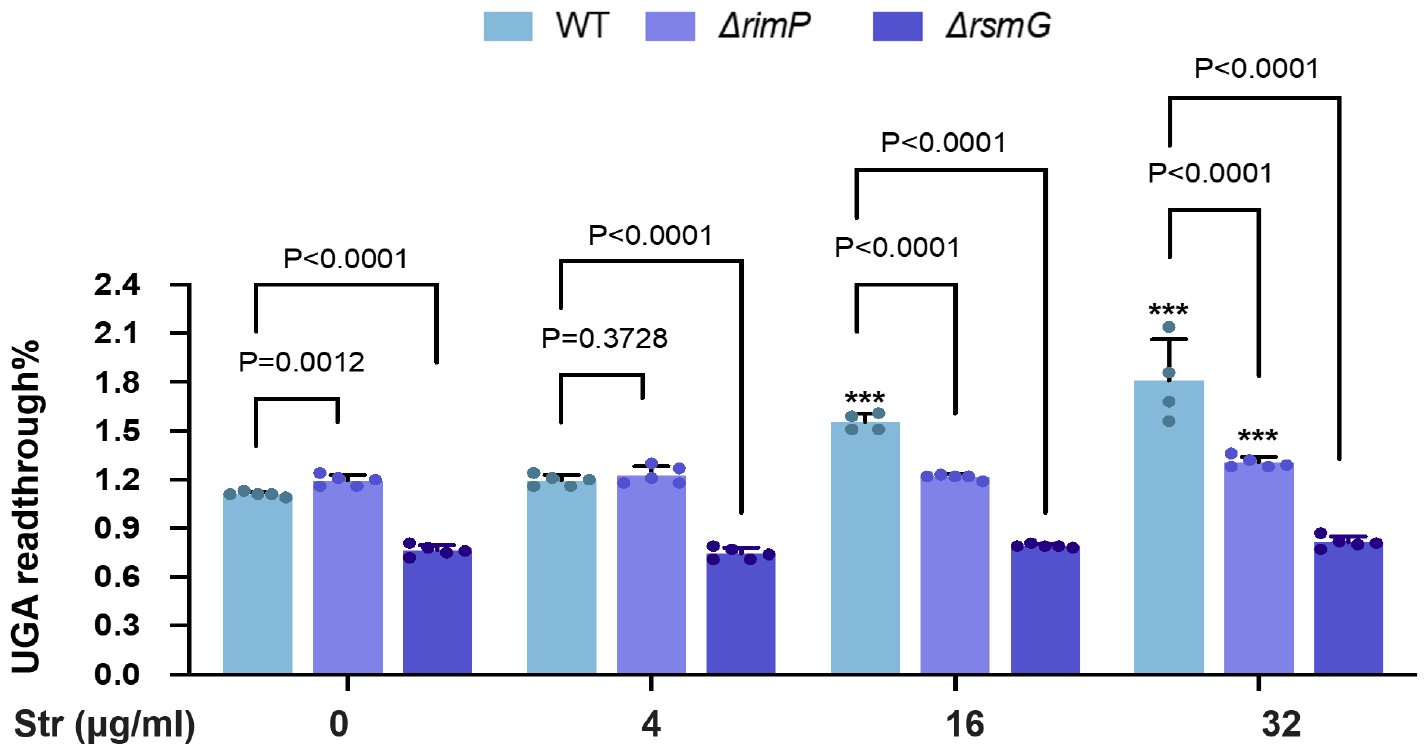
UGA readthrough of *Salmonella* strains. The UGA readthrough of stationary phase *Salmonella* carrying fluorescence reporters grown in LB with or without Str was calculated as described (36). Error bars represent one SD from the mean. The statistical differences were analyzed using One-way ANOVA with Dunnett correction. *** P < 0.001 comparing the same strain in the presence and absence of Str.

### RimP is critical for *Salmonella* motility

Antibiotic resistance is frequently accompanied by a cost of fitness (37). Our previous work shows that some mutations affecting ribosomal fidelity impair flagellar motility (38, 39), prompting us to investigate the motility of Str-resistant *Salmonella* mutants. A soft-agar swimming motility assay shows that eight out of nine evolved Str-resistant strains and the Δ*rimP* mutant are defective in flagellar motility (Fig. 5 and Table 1). To probe the expression of flagellar genes, we used a pZS P*tet*-mCherry P*fliA-*YFP reporter plasmid in platereader and fluorescence microscopy assays. FliA is a sigma factor that controls the expression of class 3 flagellar genes, and the promotor of *fliA* is directly regulated by the master flagellar regulator FlhDC (40). Consistent with the motility defect, the P*fliA* activity was substantially decreased upon deletion of *rimP*, suggesting that proper ribosome biogenesis is required for optimal expression of flagellar genes and the motility.

**Figure 5.**
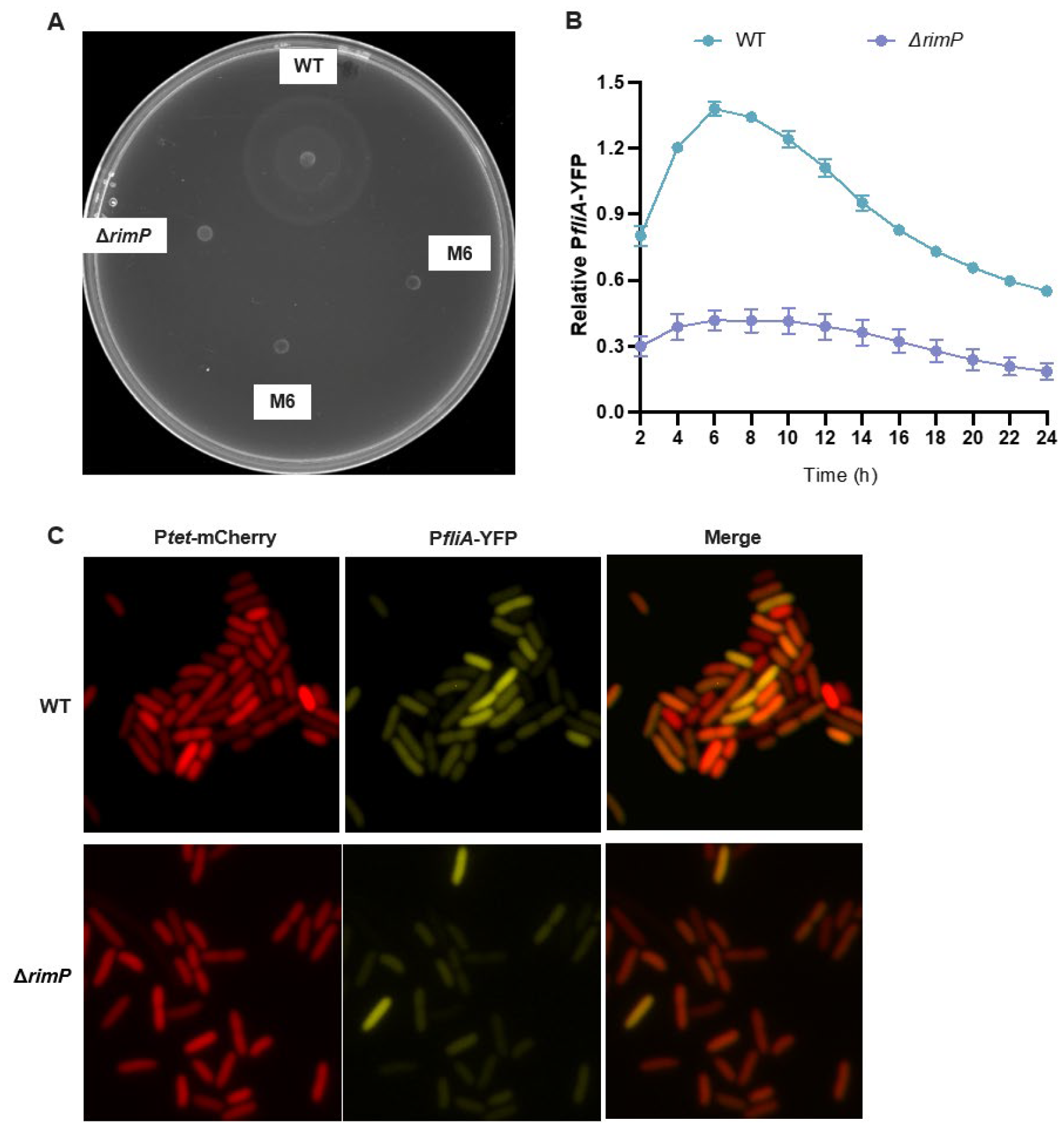
Motility of WT *Salmonella* and *rimP* mutants. (A) Soft-agar motility assay. The evolved M6 strain carrying a *rimP* frameshift mutation and the Δ*rimP* strain are both defective in flagellar motility. (B, C) Expression of P*fliA* in WT and Δ*rimP* determined using the platereader or fluorescence microscopy, respectively. P*tet*-mCherry is constitutively expressed and used for normalization. (A) and (C) show representative images of at least three biological replicates. Error bars in (B) represent one SD from the mean.

### Ribosome biogenesis affects Str resistance in *E. coli*

RimP is a conserved bacterial protein involved in the early-stage assembly of the 30S subunit (23, 41). As in *Salmonella*, deleting *rimP* also increased Str resistance in *E. coli* (Fig. 6). To assess how other ribosome biogenesis factors affect Str resistance, we tested *E. coli* mutants that lack *rbfA* (30S assembly), *rimM* (30S assembly), *rhlE* (50S assembly), *ksgA* (methylation of 16S rRNA), *rsgA* (30S-dependent GTPase), or *srmB* (50S assembly) genes. In addition to *rimP*, deleting *rhlE, ksgA, rsgA*, or *srmB* also increased resistance to Str, whereas deleting *rbfA* or *rimM* had no effect on Str resistance. These results suggest that multiple, but not all, ribosome biogenesis defects impact resistance to Str.

**Figure 6.**
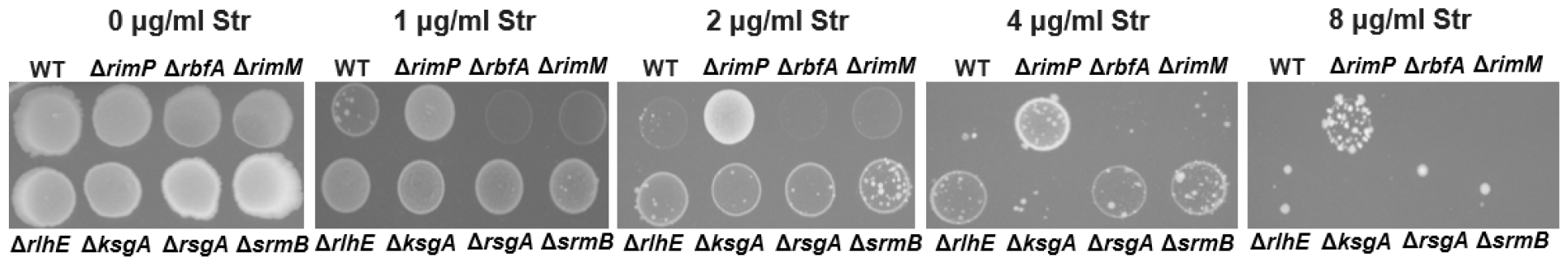
Growth of WT *E. coli* (BW25113) and mutants defective in ribosome biogenesis in the presence and absence of Str. The images are representatives of at least three biological replicates.

## Discussion

Bacteria develop antibiotic resistance through either the acquisition of foreign resistance genes or mutations of endogenous genes (42). The common resistance mechanisms include (a) prevention of access to target, including decreased uptake and increased efflux, (b) mutation or modification of the antibiotic target, and (c) modification and inactivation of the antibiotic (42, 43). All of the above mechanisms have been shown to result in resistance to aminoglycosides, which bind to the ribosome and cause protein mistranslation (42, 44). Unlike other ribosome-targeting antibiotics that are bacteriostatic, aminoglycosides are bactericidal due to their mistranslation-promoting property (4, 9). A recent study shows that aminoglycosides induce clusters of translational errors following the first mistranslation event (9), which promotes protein misfolding and aggregation (11). Aminoglycoside uptake occurs through two energy-dependent phases (EDP) EDPI and EDPII (7). The slow uptake during EDPI requires the proton motive force (PMF) driven by the membrane potential (ΔΨ) and the proton gradient (ΔpH). The fast uptake during EPDII follows EPDI and depends on the action of aminoglycosides on the ribosome. The commonly accepted model is that aminoglycosides first enter the cytoplasm in EDPI and cause mistranslation of membrane proteins, resulting in membrane leakage and faster uptake in EPDII (6, 7). In the absence of Str, the Δ*rimP*, Δ*rsmG*, and *rpsL\** mutants exhibit similar ΔΨ as the WT, as shown by the fluorescence signals of DiBAC_4_(3) (Fig. 3C), implying that EDPI is not affected by these mutations. However, in the presence of Str, all three mutants display lower entry of DiBAC_4_(3) into the cells, suggesting less membrane leakage. We also show that deleting *rimP* or *rsmG* decreased stop-codon readthrough errors in the presence of Str (Fig. 4). Collectively, our data suggest that RimP and RsmG are critical for sensitizing ribosomes to Str binding and promoting protein mistranslation.

RsmG (previously known as GidB) is responsible for the m^7^G527 modification in the 530 loop of 16S rRNA where Str binds (21, 22). Mutations in *rsmG* have been shown to increase translational fidelity in *M. tuberculosis* and *E. coli* (27, 34, 45) and cause Str resistance in *M. tuberculosis, E. coli*, and *S. enterica* (18, 20, 21, 27). It is likely that ribosomes lacking the m^7^G527 modification bind Str with a lower affinity. How ribosome assembly factors (e.g., RimP) affect aminoglycoside binding remains unclear. Cryo-electron microscopy structures of the 30S ribosomal subunit show that RimP interacts with KsgA and delays h44 to recruit uS12 in the decoding center, allowing the 30S to properly mature before assembly into the 70S ribosome (24). Interestingly, deleting *ksgA* also increases Str resistance in *E. coli* (Fig. 6). During the assembly of the ribosome, kinetic traps can lead to alternative rRNA conformations (46). It is possible that the 70S ribosomes assembled in the absence of RimP or KsgA may adopt a conformation in the decoding center that decreases the affinity for Str. It would be intriguing to test this model using structural and biophysical analyses in future studies.

## Materials and Methods

### Bacterial strains, plasmids, growth conditions, and reagents

All *Salmonella* strains used in this study are derived from *S*. Typhimurium ATCC 14028s. Bacterial strains and plasmids are listed in Table S1. Gene deletion mutants were constructed by lambda red-mediated recombination as previously described (47), and the oligonucleotides used are listed in Table S2. Unless otherwise noted, all bacteria used in this study were grown in Luria-Bertani (LB) Lennox media containing 10 g/l tryptone, 5 g/l yeast extract, and 5 g/l NaCl. Antibiotics were added when needed (100 μg/ml ampicillin and 25 μg/ml chloramphenicol).

### Laboratory evolution of Str-resistant strains

Briefly, 9 independent populations of *S*. Typhimurium ATCC 14028s were propagated in 96-well microtiter plates containing 100 μl LB media with Str concentrations ranging from 0.5 to 128 μg/ml. 2 μl of bacterial culture were transferred every 24 h to fresh LB with increasing concentrations of Str. The experiment halted when the evolving populations reached an up to 16-fold increase in resistance relative to the WT ancestor. The isolated clones (M1-M9) were frozen in glycerol stocks at −80°C for further whole-genome sequencing.

### Whole-genome sequencing and identification of mutations

PE150 libraries of the *Salmonella* strains were prepared and sequenced by Novogene. Reads are deposited in Sequence Read Archive (SRA) under accession number SUB14099288. Reads were trimmed using TrimGalore! and mapped to the genome of *S. typhimurium* ATCC 14028 (assembly ASM325338v1) using Bowtie2. To identify mutations, we used FreeBayes (https://arxiv.org/abs/1207.3907) for the detection of candidate variants. The Integrated Genome Viewer (https://software.broadinstitute.org/software/igv/download) was used to inspect candidate variants. True mutations were differentiated from sequencing errors and preexisting SNPs by being supported by the consensus of the reads in the evolved isolate(s), but not by the reads from the starting WT strain.

### Growth rate measurement

Overnight cultures were diluted 1:100 in fresh LB in quadruplicates. The cultures were incubated at 37°C for 16 h with shaking, and A_600_ measurements were taken every 20 min using a platereader (Synergy HTX, BioTek). The growth rates of the exponential phase were calculated.

### MIC assay

The *Salmonella* strains were grown overnight in LB medium at 37 °C with shaking. The next day, all cultures were normalized to A_600_ ∼2.0, and diluted 1:100 in fresh LB. The MICs for different antibiotics were determined by spotting 5 μL of diluted cultures onto LB agar plates with 2-fold increasing concentrations of Str, tetracycline, ciprofloxacin, or ampicillin. The plates were incubated at 37 °C for 1.5 days, and the MICs were set to the lowest concentration of antibiotic with no visible bacterial lawn formation.

### Spontaneous mutation rates

The rates of mutations leading to high-Str resistance were estimated using fluctuation tests as described (48). Briefly, the *Salmonella* strains were grown at 37°C with continuous shaking. The next day, the normalized cultures were diluted 1:100 in fresh LB and grown aerobically for 2∼3 h at 37°C until A_600_ reached 0.2–0.4. The resulting cultures were then diluted in LB to approximately 5,000 cells per ml, transferred into 96-well microtiter plates, and incubated for 24 h at 37°C. The cultures of each well were individually plated on LB agar supplemented with 512 μg/ml Str to determine the number of spontaneous mutants that arose during growth. In parallel, the diluted cultures were plated on LB agar to determine the total viable cells. The mutation rate of each tested strain was estimated using the number of Str-resistant mutants divided by the total number of cells.

### Efflux activity assay

Overnight cultures were diluted 1:100 into fresh LB and incubated for 2∼3 h at 37°C. All cultures were normalized to A_600_ ∼0.5, and 100 μl aliquots were transferred to black polystyrene 96-well plates. The cells were incubated with 50 μg/ml Nile red with agitation for 2∼3 h at 37°C. The fluorescence intensity was measured in a platereader (Synergy H1, BioTek) using an excitation wavelength of 549 nm and an emission wavelength of 628 nm.

### Membrane potential measurement

Overnight cultures were diluted 1:100 into fresh LB and incubated for 2∼3 h at 37°C. All cultures were normalized to A_600_ ∼0.5 and treated with or without 128 or 1024 μg/ml Str. DiBAC_4_(3) dissolved in dimethyl sulfoxide (DMSO) was then added to a final concentration of 5 μM, and the fluorescence intensity was monitored every 20 min in a platereader (Synergy H1, BioTek) for 18 h using an excitation wavelength of 493 nm and an emission wavelength of 516 nm. Single-cell fluorescence was measured using the FACSCanto II flow cytometer at a low flow rate. 30,000 events were collected for each sample, and the FlowJo software was used for further data analysis.

### Streptomycin uptake assay

Overnight cultures were diluted 1:100 into fresh LB and incubated for 2∼3 h at 37°C. All cultures were normalized to A_600_ ∼0.5 and supplemented with 1024 μg/ml Str. Following 3 h of incubation at 37°C with agitation, 1 ml of each culture was harvested by centrifugation, washed three times with phosphate-buffered saline (PBS), and finally resuspended into 300 μl PBS. For estimation of Str uptake, bacterial cells were lysed by sonication and 5 μl of the culture supernatants were spotted on agar plates with an *E. coli* (MG1655) lawn. Plates were incubated at 37 °C for 1.5 days, and the zones of growth inhibition were compared between WT and the Δ*rimP* mutant.

### Fluorescence-based stop-codon readthrough assay

Stop-codon readthrough rates with or without Str were determined using plasmids pZS-P*tet*-m-TGA-y, pZS-P*tet*-m-y, and pZS-P*tet*-*lacZ* as described (36).

### Soft-agar motility assay

Overnight cultures were diluted 1:100 into fresh LB and incubated for 2∼3 h at 37°C. All cultures were normalized to A_600_ ∼0.5. 3 μl of each culture was spotted on LB plates with 0.25% agar, and incubated for 6 h at 37 °C. The diameters of the bacterial rings were measured, and the images were taken in the ChemiDoc Imaging System (Bio-Rad).

### Measurement of promoter activity with a platereader

Overnight cultures were diluted 1:100 and grown in LB Miller containing 10 g/l tryptone, 5 g/l yeast extract, and 10 g/l NaCl at 37°C with vigorous shaking. The fluorescence was measured every 20 min using a platereader (Synergy HTX, BioTek) for 24 h under the optimal excitation and emission wavelengths for each fluorescence protein (Ex = 575 nm and Em = 620 nm for mCherry; Ex = 508 nm and Em = 560 nm for YFP). The gain was set to 40. The P*filA* promoter activity was calculated as the ratio of YFP over mCherry.

### Fluorescence microscopy

Overnight cultures carrying the pZS P*tet*-mCherry P*fliA*-YFP P*fliC*-eCFP plasmid were diluted 1:50 in LB, and Cells were grown aerobically in LB Miller at 37°C for 5 h to the early stationary phase prior to imaging. 1 μl of each culture was placed on a 1.5% agarose LB pad on a 12-well slide. Fluorescence images were obtained with a 100 X oil lens using BZ-X800 fluorescence microscope (Keyence). Image analysis was performed using ImageJ/Fiji (NIH).

## Statistical analysis

The data presented corresponds to the mean with standard deviation (SD) values of at least three biological replicates. The statistical differences were analyzed using One-way ANOVA with Dunnett correction, where P < 0.05 was considered significant (*) and P < 0.001 as highly significant (***).

## Data Availability

Genome sequencing reads are deposited in Sequence Read Archive (SRA) under accession number SUB14099288.

## Supplementary Materials

Tables S1-S2 and Figure S1-S2.

## Acknowledgments

This work was funded by the National Institute of General Medical Sciences (R35GM136213 to J.L. and R35GM141710 to A.v.H.).

## Author Contributions

Z.L., Y.L., and J.L. designed the project; Z.L. and Y.L. performed the experiments; Z.L., Y.L., A.v.H., and J.L. analyzed the data and wrote the manuscript.

## References

1. Antimicrobial Resistance C. 2022. Global burden of bacterial antimicrobial resistance in 2019: a systematic analysis. Lancet 399:629–655.

2. Bottger EC, Crich D. 2020. Aminoglycosides: time for the resurrection of a neglected class of antibacterials? ACS Infect Dis 6:168–172.

3. Anonymous. 2018. Critically important antimicrobials for human medicine. World Health Organization:6th revision.

4. Wilson DN. 2014. Ribosome-targeting antibiotics and mechanisms of bacterial resistance. Nat Rev Microbiol 12:35–48.

5. Lin J, Zhou D, Steitz TA, Polikanov YS, Gagnon MG. 2018. Ribosome-targeting antibiotics: modes of action, mechanisms of resistance, and implications for drug design. Annu Rev Biochem 87:451–478.

6. Davis BD. 1987. Mechanism of bactericidal action of aminoglycosides. Microbiol Rev 51:341–50.

7. Lang M, Carvalho A, Baharoglu Z, Mazel D. 2023. Aminoglycoside uptake, stress, and potentiation in Gram-negative bacteria: new therapies with old molecules. Microbiol Mol Biol Rev doi:10.1128/mmbr.00036-22:e0003622.

8. Schatz A, Bugie E, Waksman SA. 2005. Streptomycin, a substance exhibiting antibiotic activity against gram-positive and gram-negative bacteria. 1944. Clin Orthop Relat Res doi:10.1097/01.blo.0000175887.98112.fe:3-6.

9. Wohlgemuth I, Garofalo R, Samatova E, Gunenc AN, Lenz C, Urlaub H, Rodnina MV. 2021. Translation error clusters induced by aminoglycoside antibiotics. Nat Commun 12:1830.

10. Vallabhaneni H, Farabaugh PJ. 2009. Accuracy modulating mutations of the ribosomal protein S4-S5 interface do not necessarily destabilize the rps4-rps5 protein-protein interaction. Rna 15:1100–9.

11. Ling J, Cho C, Guo LT, Aerni HR, Rinehart J, Söll D. 2012. Protein aggregation caused by aminoglycoside action is prevented by a hydrogen peroxide scavenger. Mol Cell 48:713–22.

12. Davis BD, Chen LL, Tai PC. 1986. Misread protein creates membrane channels: an essential step in the bactericidal action of aminoglycosides. Proc Natl Acad Sci U S A 83:6164–8.

13. Carter AP, Clemons WM, Brodersen DE, Morgan-Warren RJ, Wimberly BT, Ramakrishnan V. 2000. Functional insights from the structure of the 30S ribosomal subunit and its interactions with antibiotics. Nature 407:340–8.

14. Demirci H, Murphy Ft, Murphy E, Gregory ST, Dahlberg AE, Jogl G. 2013. A structural basis for streptomycin-induced misreading of the genetic code. Nat Commun 4:1355.

15. Cohen KA, Stott KE, Munsamy V, Manson AL, Earl AM, Pym AS. 2020. Evidence for expanding the role of streptomycin in the management of drug-resistant Mycobacterium tuberculosis. Antimicrob Agents Chemother 64.

16. Nair J, Rouse DA, Bai GH, Morris SL. 1993. The rpsL gene and streptomycin resistance in single and multiple drug-resistant strains of Mycobacterium tuberculosis. Mol Microbiol 10:521–7.

17. Allen PN, Noller HF. 1991. A single base substitution in 16S ribosomal RNA suppresses streptomycin dependence and increases the frequency of translational errors. Cell 66:141–8.

18. Mikheil DM, Shippy DC, Eakley NM, Okwumabua OE, Fadl AA. 2012. Deletion of gene encoding methyltransferase (gidB) confers high-level antimicrobial resistance in Salmonella. J Antibiot (Tokyo) 65:185–92.

19. Dai R, He J, Zha X, Wang Y, Zhang X, Gao H, Yang X, Li J, Xin Y, Wang Y, Li S, Jin J, Zhang Q, Bai J, Peng Y, Wu H, Zhang Q, Wei B, Xu J, Li W. 2021. A novel mechanism of streptomycin resistance in Yersinia pestis: Mutation in the rpsL gene. PLoS Negl Trop Dis 15:e0009324.

20. Wistrand-Yuen E, Knopp M, Hjort K, Koskiniemi S, Berg OG, Andersson DI. 2018. Evolution of high-level resistance during low-level antibiotic exposure. Nat Commun 9:1599.

21. Okamoto S, Tamaru A, Nakajima C, Nishimura K, Tanaka Y, Tokuyama S, Suzuki Y, Ochi K. 2007. Loss of a conserved 7-methylguanosine modification in 16S rRNA confers low-level streptomycin resistance in bacteria. Mol Microbiol 63:1096–106.

22. Watson ZL, Ward FR, Meheust R, Ad O, Schepartz A, Banfield JF, Cate JH. 2020. Structure of the bacterial ribosome at 2 A resolution. Elife 9.

23. Maksimova E, Kravchenko O, Korepanov A, Stolboushkina E. 2022. Protein assistants of small ribosomal subunit biogenesis in bacteria. Microorganisms 10.

24. Schedlbauer A, Iturrioz I, Ochoa-Lizarralde B, Diercks T, Lopez-Alonso JP, Lavin JL, Kaminishi T, Capuni R, Dhimole N, de Astigarraga E, Gil-Carton D, Fucini P, Connell SR. 2021. A conserved rRNA switch is central to decoding site maturation on the small ribosomal subunit. Sci Adv 7.

25. Baym M, Stone LK, Kishony R. 2016. Multidrug evolutionary strategies to reverse antibiotic resistance. Science 351:aad3292.

26. Pezzella C, Ricci A, DiGiannatale E, Luzzi I, Carattoli A. 2004. Tetracycline and streptomycin resistance genes, transposons, and plasmids in Salmonella enterica isolates from animals in Italy. Antimicrob Agents Chemother 48:903–8.

27. Wong SY, Javid B, Addepalli B, Piszczek G, Strader MB, Limbach PA, Barry CE, 3rd. 2013. Functional role of methylation of G518 of the 16S rRNA 530 loop by GidB in Mycobacterium tuberculosis. Antimicrob Agents Chemother 57:6311–8.

28. Goltermann L, Good L, Bentin T. 2013. Chaperonins fight aminoglycoside-induced protein misfolding and promote short-term tolerance in Escherichia coli. J Biol Chem 288:10483–9.

29. Verstraeten N, Knapen WJ, Kint CI, Liebens V, Van den Bergh B, Dewachter L, Michiels JE, Fu Q, David CC, Fierro AC, Marchal K, Beirlant J, Versees W, Hofkens J, Jansen M, Fauvart M, Michiels J. 2015. Obg and membrane depolarization are part of a microbial bet-hedging strategy that leads to antibiotic tolerance. Mol Cell 59:9–21.

30. Lv B, Huang X, Lijia C, Ma Y, Bian M, Li Z, Duan J, Zhou F, Yang B, Qie X, Song Y, Wood TK, Fu X. 2023. Heat shock potentiates aminoglycosides against gram-negative bacteria by enhancing antibiotic uptake, protein aggregation, and ROS. Proc Natl Acad Sci U S A 120:e2217254120.

31. Koronakis V, Eswaran J, Hughes C. 2004. Structure and function of TolC: the bacterial exit duct for proteins and drugs. Annu Rev Biochem 73:467–89.

32. Bjorkman J, Samuelsson P, Andersson DI, Hughes D. 1999. Novel ribosomal mutations affecting translational accuracy, antibiotic resistance and virulence of Salmonella Typhimurium. Mol Microbiol 31:53–8.

33. Agarwal D, Gregory ST, O’Connor M. 2011. Error-prone and error-restrictive mutations affecting ribosomal protein S12. J Mol Biol 410:1–9.

34. Lyu Z, Villanueva P, O’Malley L, Murphy P, Augenstreich J, Briken V, Singh A, Ling J. 2023. Genome-wide screening reveals metabolic regulation of stop-codon readthrough by cyclic AMP. Nucleic Acids Res doi:10.1093/nar/gkad725.

35. Fan Y, Thompson L, Lyu Z, Cameron TA, De Lay NR, Krachler AM, Ling J. 2019. Optimal translational fidelity is critical for Salmonella virulence and host interactions. Nucleic Acids Res 47:5356–5367.

36. Fan Y, Evans CR, Barber KW, Banerjee K, Weiss KJ, Margolin W, Igoshin OA, Rinehart J, Ling J. 2017. Heterogeneity of stop codon readthrough in single bacterial cells and implications for population fitness. Mol Cell 67:826–836.

37. Andersson DI, Hughes D. 2010. Antibiotic resistance and its cost: is it possible to reverse resistance? Nat Rev Microbiol 8:260–71.

38. Fan Y, Evans CR, Ling J. 2016. Reduced protein synthesis fidelity inhibits flagellar biosynthesis and motility. Sci Rep 6:30960.

39. Lyu Z, Yang A, Villanueva P, Singh A, Ling J. 2021. Heterogeneous flagellar expression in single Salmonella cells promotes diversity in antibiotic tolerance. mBio 12:e0237421.

40. Chilcott GS, Hughes KT. 2000. Coupling of flagellar gene expression to flagellar assembly in Salmonella enterica serovar Typhimurium and Escherichia coli. Microbiol Mol Biol Rev 64:694–708.

41. Shajani Z, Sykes MT, Williamson JR. 2011. Assembly of bacterial ribosomes. Annu Rev Biochem 80:501–26.

42. Darby EM, Trampari E, Siasat P, Gaya MS, Alav I, Webber MA, Blair JMA. 2023. Molecular mechanisms of antibiotic resistance revisited. Nat Rev Microbiol 21:280–295.

43. Blair JM, Webber MA, Baylay AJ, Ogbolu DO, Piddock LJ. 2015. Molecular mechanisms of antibiotic resistance. Nat Rev Microbiol 13:42–51.

44. Ramirez MS, Tolmasky ME. 2010. Aminoglycoside modifying enzymes. Drug Resist Updat 13:151–71.

45. Bi Z, Su HW, Hong JY, Javid B. 2021. Ribosomal RNA methylation by GidB is a capacitor for discrimination of mischarged tRNA. bioRxiv:2021.03.02.433644.

46. Woodson SA. 2000. Recent insights on RNA folding mechanisms from catalytic RNA. Cell Mol Life Sci 57:796–808.

47. Datsenko KA, Wanner BL. 2000. One-step inactivation of chromosomal genes in Escherichia coli K-12 using PCR products. Proc Natl Acad Sci U S A 97:6640–5.

48. Swings T, Van den Bergh B, Wuyts S, Oeyen E, Voordeckers K, Verstrepen KJ, Fauvart M, Verstraeten N, Michiels J. 2017. Adaptive tuning of mutation rates allows fast response to lethal stress in Escherichia coli. Elife 6.

